# Male and female syringeal muscles exhibit superfast shortening velocities in Zebra finches

**DOI:** 10.1101/2023.06.19.545584

**Authors:** Nicholas W. Gladman, Coen P. H. Elemans

**Affiliations:** Vocal Neuromechanics lab, Sound Communication and Behaviour group, Department of Biology, University of Southern Denmark, 5230 Odense M, Denmark

**Keywords:** Muscle performance, isovelocity, Vmax, vocal performance, song learning

## Abstract

Vocalisations play a key role in the communication behaviour of many vertebrates. Vocal production requires extremely precise motor control, which is executed by superfast vocal muscles that can operate at cycle frequencies over 100 Hz and up to 250 Hz. The mechanical performance of these muscles has been quantified with isometric performance and the workloop technique, but due to methodological limitations we lack a key muscle property characterising muscle performance, the force-velocity (FV) relationship. Here we establish a method that allows quantification of the FV relationship in extremely fast muscles, and test if the maximal shortening velocity of zebra finch syringeal muscles is different between males and females. We show that syringeal muscles exhibit extremely high maximal shortening velocities of 46 L_0_ s^-1^, far exceeding most other vocal and skeletal muscles, and that isometric properties positively correlate with maximal shortening velocities. While male and female muscles differ in isometric speed measures, maximal shortening velocity surprisingly is not sex-dependent. We also show that cyclical methods to measure force-length properties used in classical laryngeal studies give the same result as conventional stepwise methodologies, suggesting either approach is appropriate. Next to force, instantaneous power also trades for speed, further highlighting these muscles are tuned to operate at high frequencies. We argue that the high thermal dependence of superfast vocal muscle performance may impact vocal behaviour.

**Summary statement:** Zebra finch syringeal muscle exhibits superfast shortening of 46 L_0_ s^-1^. Shortening is not sex-specific but correlates with isometric performance – faster twitches and tetani are associated with faster shortening.

## Introduction

Vocal communication is of key importance to vertebrates, with vocalisations playing important roles in behaviours ranging from predator avoidance, through to conspecific recognition, and mate choice (Bradbury and Vehrencamp, 2011; Kelley et al., 2008; Nowicki and Searcy, 2004). Vocal signalling demands a high degree of motor control precision, and the premotor circuitry in songbirds and mammals indeed has millisecond precision (Chi and Margoliash, 2001; Nieder and Mooney, 2020; Pomberger et al., 2018). The precise execution of descending motor control is facilitated by the vocal muscles of the mammalian larynx and avian syrinx that, by modifying vocal fold tension and positioning, control key acoustic parameters such as amplitude and fundamental frequency (Riede and Goller, 2010). In both the larynx and syrinx superfast muscles have evolved that are optimised for speed (Elemans et al., 2008; Elemans et al., 2011). Superfast muscles produce positive work ≥100 cycles/sec and those in larynx and syrinx can modulate vocal features up to 200-250 Hz (Elemans et al., 2008; Mead et al., 2017). To achieve such remarkable performance, superfast muscles exhibit several cellular and molecular adaptations, such as large volume-proportions of sarcoplasmic reticula, amplified expression of calcium-handling systems, increased fibre diameter, as well as expressing unique myosin isoforms (Hoh, 2005; Mead et al., 2017; Rome, 2006; Rome and Lindstedt, 1998), that allow them to operate at the maximum operational speed set by fundamental constraints in synchronous muscle architecture (Mead et al., 2017). These adaptations for speed however come at the expense of force, because of reduced myofibrillar area and lower number of formed cross bridges (Rome and Lindstedt, 1998; Rome et al., 1996). Superfast muscles evolved independently in sound-producing organs in ray-finned fish, birds, and mammals, suggesting a paramount importance in vocal communication (Mead et al., 2017).

The mechanical performance of muscle depends on both length and the velocity of shortening. Both force-length and force-velocity relationships aid to predict *in vivo* mechanical behaviour of muscles. However, the assessment of the mechanical properties in avian vocal muscles has largely focused on isometric (Adam et al., 2023) and cyclic performance (Elemans et al., 2008; Elemans et al., 2006), but not the force-velocity relationship. Force-velocity properties have previously been measured in mammalian laryngeal muscles and amphibian vocal muscles, such as those of the baboon (Mardini et al., 1987), rat (McMullen and Andrade, 2006), human (D’Antona et al., 2002; Sciote et al., 2002), dog (Alipour and Titze, 1999; Toniolo et al., 2007), and the hylid frogs (Girgenrath and Marsh, 2003; McLister et al., 1995), that are all slower than avian superfast syrinx muscles. Attempts to measure the force-velocity relationship in superfast avian syringeal muscle have been hampered by the low force generating capacity and extremely fast contraction times of these muscles. Quick release methodologies have proved ineffective due to muscle recovering force too rapidly, while force-clamp methodologies could not be effectively employed due to the low force generating ability of this muscle. We thus lack insight in a critical feature describing the mechanical performance of superfast syringeal muscles.

While vocal communication is exhibited by both sexes across lineages, the role played in mate selection has led to sexual dimorphism of this trait in some species. Sexual dimorphism is common in songbirds, where birds such as the zebra finch (*Taeniopygia guttata*), show clear male-female differences. Male zebra finches use song to attract mates and enhance pair-bonding (Griffith, 2019; Loning et al., 2022), while females do not produce song. Females have considerably smaller syringeal muscles than males (Christensen et al., 2017; Wade and Buhlman, 2000). Alongside these anatomical differences, functional differences of the muscles themselves are also present. Elemans *et al*. (2008) demonstrated male syringeal muscles isometric properties are nearly twice as fast as female syringeal muscles (Adam and Elemans, 2020; Elemans et al., 2008), and express more myosin heavy-chain gene MYH13 with superfast properties (Mead et al., 2017). However, we currently lack non-isometric speed measures in female muscles.

The key aim of this study is to establish a technique to measure the force-velocity properties of the superfast syringeal muscles and test if male and female zebra finch muscles perform differently. Due to their higher isometric contraction speeds, we hypothesise that male syringeal muscles have higher shortening velocities than females.

To enhance our understanding of vocal muscle performance, we furthermore assessed two methodologies of assessing force-length profiles. Alipore-Haghighi (1991) utilised a technique where laryngeal muscle was subject to cyclical length changes at 1 Hz both passively and actively to find force-length properties of the muscle. Such long stimulations potentially deplete the energy storage and fatigue the muscle and therefore are not routinely used in conventional muscle physiology. Here we assessed if this methodology results in similar force-length profiles and optimal length (*L*_*0*_) measured as commonly used stepwise approaches. The cyclic methodology involves maximally stimulating a muscle while undergoing a length change, this methodology does not allow the muscle to relax between successive stimuli, which likely impacts cross-bridge formation, breaking, as well as causing residual passive force enhancement due to the action of titin and other non-contractile elements (Herzog, 2019). We therefore tested both methods to assess its suitability for testing this key muscle performance parameter.

## Methods

We settled on using the isovelocity technique to assess force-velocity properties of syringeal muscle. This technique involves controlling muscle shortening velocity rather than force, where the muscle is shortened at set velocities and the resulting force measured.

### Bird housing conditions

Adult zebra finches (*Taeniopygia guttata*, Vieillot (1817)) were housed within an aviary at the University of Southern Denmark. Animals were kept in a mixed-sex group of approximately 100 animals, on a 13:11h light cycle. Food, water, and cuttlebones (*Sepia* spp.) were available to all animals ad-libitum within the aviary. We used 6 female and 8 male zebra finches, at the time of use female animals were 182 ± 70 days old, and males were 339 ± 142 days old.

### Muscle fascicle preparation and mounting

Zebra finches were euthanised in accordance with Danish law, as approved by the Danish Animal Experiments Inspectorate (Copenhagen, Denmark), *via* an overdose of isoflurane (Attane vet, ScanVet Animal Health A/S, Fredensborg, Denmark). The syrinx was extracted through a vertical incision along the sternum, during the dissection process the chest cavity was regularly flushed with ice-cold oxygenated avian ringers solution (see supplementary table 1 for composition; Adam *et al*., 2021). The extracted syrinx was placed in 1 ± 1 °C oxygenated avian ringers solution within a petri-dish, temperature was maintained using an aluminium cooling plate with constant circulation through a chiller (Lauda RM6 Refrigerated Circulating Bath, Lauda Dr. R. Wobser GMBH & co., Lauda, Germany).

We focussed on the dorsal tracheobronchial muscle (DTB) where we have most information on isometric contractile performance (Adam and Elemans, 2019; Adam and Elemans, 2020; Adam et al., 2021; Adam et al., 2023; Elemans et al., 2008; Mead et al., 2017). The DTB was dissected out of the extracted syrinx. The side from which muscle was dissected out was randomly determined to minimise left-right biases (8/14 57% from left, 6/14 43% from right). A small piece of bone (bronchial half-ring B2) and associated connective tissue (clavicular air sac membrane, CASM) were kept at either end of the muscle. We securely fastened both ends into aluminium foil T-clips (Photofabrication Ltd, St Neots, Cambs, UK). Clips were used to connect the muscle to 0.1 mm stainless steel hooks (made from Austerlitz Insect pins Minutiens 0.1 mm, Entomoravia, Slavkov u Brna, The Czech Republic) connected to a force-transducer (400A, Aurora Scientific inc., Aurora, ON, Canada) at one end, and a high-speed length controller (322C, Aurora Scientific inc.) at the other. Both the length controller and force transducer were mounted on 3D micro-manipulators. The muscle was held within a temperature-controlled bath (cooled *via* an 826A system, Aurora Scientific inc.) and continually superfused with ringers solution using a peristaltic pump (Ole Dich Instrumentmakers ApS, Hvidovre, Denmark). Platinum electrodes ran parallel to the muscle on both sides and were used for electrical stimulation of the muscle *via* a high-power, bi-phase stimulator (701C Aurora Scientific inc.). Signals were low pass filtered (10 kHz, inline filter, Thor labs, Newton, NJ, USA) and digitised at 40 kHz, 16 bits (PCI 6259, National Instruments, Austin, TX, USA). All control software was written in MATLAB (2021b, Mathworks, Natick, MA, USA).

### Protocol

At a temperature of 30 ± 0.5°C, slack muscle length was measured using digital callipers, and adjusted to 110% of this initial length using a micromanipulator. The stimulation conditions were optimised using a series of tetani, this was determined by modifying pulse amplitude and duration. Next, we increased the stimulation frequency from 100 to 500 in 50 Hz steps to find the tetanic frequency where isometric force was maximal.

#### Force-Length curve

We next determined the optimal length (*L*_*0*_) of muscle by applying a series of isometric tetani at different lengths. This began at 110% slack length, with length modified in steps of 0.25 mm. Modification of muscle length began by increasing the length from the initial length. If muscle force declined, the muscle length was decreased from the initial length. The length at which maximal force was recorded was defined as *L*_*0*_. Between each successive tetani the muscle was allowed to rest for 3 minutes.

After length optimisation, the muscle was stimulated to produce an isometric twitch and tetanus at *L*_*0*_.

#### Force-velocity curves

Force-velocity curves were determined using the isovelocity technique (based on Claflin and Faulkner (1989)). Preliminary tests using other methodologies (slack tests and force-holds) proved ineffective, where muscular force would recover too rapidly during slack test measures, and muscle was too weak to reliably use force-holds in a dual-mode lever system (Aurora Scientific, model 300C). During isovelocity measures, we set the length to 110% *L*_*0*_, and tetanised the muscle (**Figure 1**). While stimulating, we shortened the muscle at constant velocity back to *L*_*0*_. Once stimulation stopped the muscle was gradually returned to 110% *L*_*0*_ at 0.3 L_0_ s^-1^. This procedure was repeated for a range of isovelocity values from 0.25 to 35 L_0_ s^-1^ in series from the fastest value down to the slowest to minimise muscle fatigue. To correct for passive force as well as the impacts of serial elasticity, we performed two runs per isovelocity value: one with stimulation (active) and one without stimulation (passive). The muscle was given 3 minutes to rest between successive active isovelocity runs.

**Figure 1:**
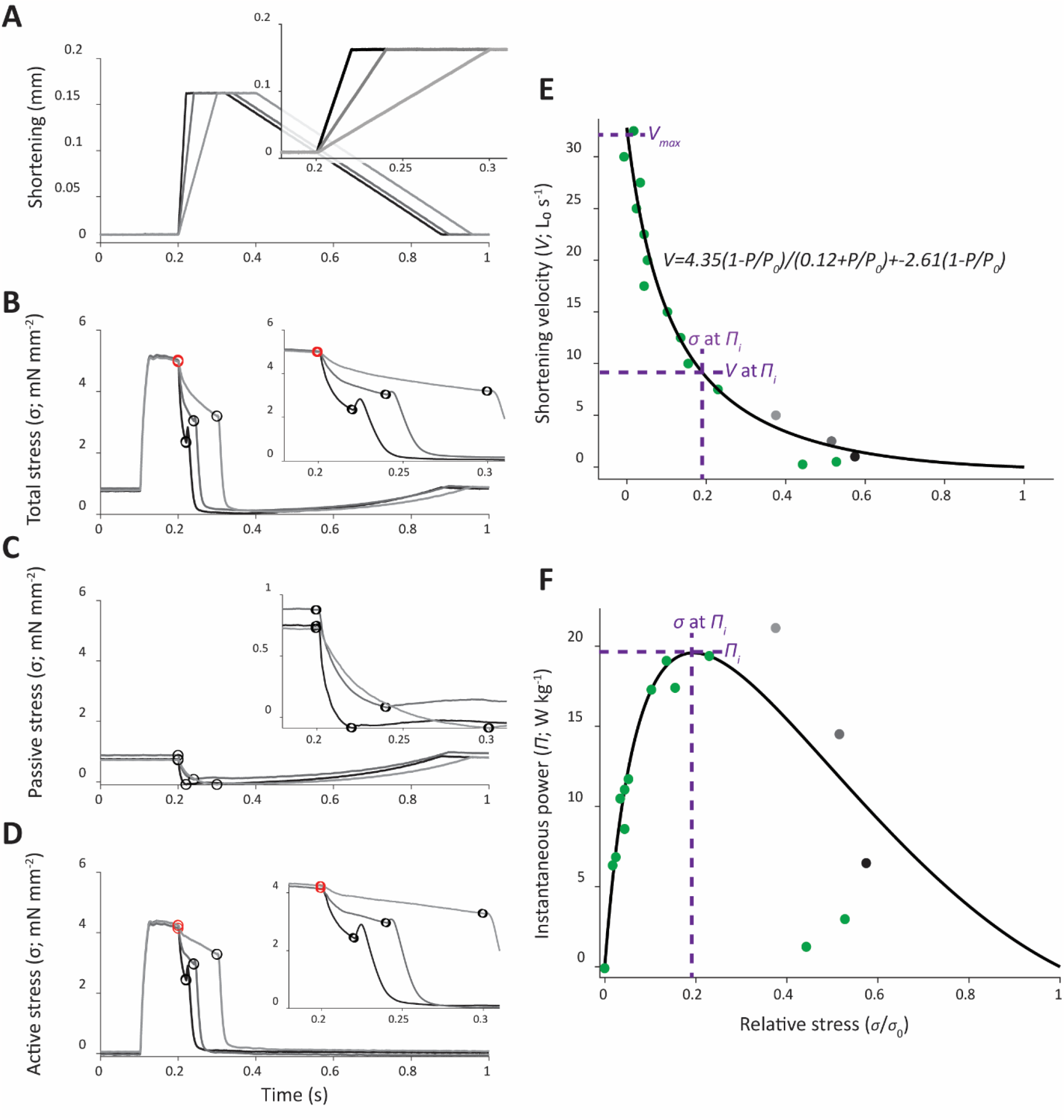
Force-velocity protocol and analysis example. (A) Length changes at different ramp velocities. (B) Total muscular stress when stimulated where the muscle tetanised before being shortened at the differing ramp velocities (C) Passive stress during shortening, (D) Active stress, calculated as (B) – (C). Inset panels in (A)-(D) show a zoomed version of the ramp process, highlighting the points from which stress measures were obtained. Graded shades of grey are used represent different ramp velocities; circles show the area where tetanic force (red) and force (black) were measured. (E) Resulting force-velocity profile including the data shown in (A)-(D) in corresponding colours. (F) Power output derived from the force-velocity profile; powers calculated from shown data are displayed in the same colours. Purple lines on (E) – (F) are used to highlight the maximum shortening velocity (*V*_*max*_), stress at peak power (*σ* at *Π*_*i*_), velocity at peak power (*V* at *Π*_*i*_) and peak power (*Π*_*i*_). Data shown is left DTB preparation from female BY155 at 30°C.

At a body temperature of approximately 40°C, the muscles shorten too rapidly for the motor of the high-speed length controller. Force-velocity curves were therefore determined at two temperatures: 20 and 30°C, where maximum shortening velocity could be measured accurately.

#### Comparison of force-length curves obtained via stepwise and cyclic methodologies

Next, we tested if stepwise and continuous cyclical protocols result in different force-length curves. To ensure muscle was not damaged during sustained stimulation, cyclic methods were carried out after force-velocity measures. At *L*_*0*_ and 30°C, muscle fascicles were subject to two 1 Hz sinusoidal length changes (± 0.5 mm from *L*_*0*_). This consisted of one unstimulated (passive) length change, and one stimulated (active) length change (following Alipour-Haghighi *et al*. 1991).

#### CSA of preparation

Lastly, we determined the length and cross-sectional area of the preparation. We placed a 5 mm silver right angle prism mirror (MRA05-P01, Thor labs) next to the preparation in the bath and took an image including the top and sideview using a camera mounted on a fluorescent stereo microscope (Flexacam C3 on a M165 FC, Leica Microsystems ltd., Heerbrugg, Switzerland). From this image we measured muscle width (*M*_*w*_) and depth (*M*_*d*_) at three points along the muscle fibre bundle using ImageJ v1.53 (Rasband, 2018). These three points were used to calculate the average width and depth of the muscle. From these, we calculated CSA assuming an ellipsoid cross-section: 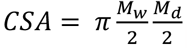.

### Data analysis

All data were analysed using MATLAB (2021b, Mathworks).

#### Force-length

At each length step, the active and passive force was obtained and used to plot force-length profiles. In, the stepwise approach passive force was the average force during the first 100 ms prior to stimulation. Active force was calculated as the average force over the plateau region of the tetanus minus the passive force. The force plateau was detected as the region between the first instance of the maximum value and the point at which this changes significantly (*findchangepts* function). These points were visually assessed to ensure only the plateau region was sampled.

The cyclic approach used the unstimulated lengthening-shortening cycle as the passive force. Active force was calculated by subtracting the passive force from the stimulated force trace at each time point. To ensure data were aligned, the start and endpoints were set and detected using the *find* function in MATLAB. All forces (*P*) were converted to stresses (*σ*) by dividing by the muscle cross-sectional area (CSA).

#### Force-velocity

The force-velocity relationship of a muscle is described by the hyperbolic-linear equation (Marsh and Bennett, 1986):

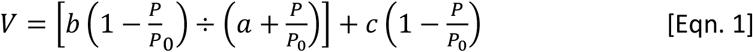

Where *a, b* and *c* represent constants, *P* is force and *P*_*0*_ is peak isometric force. Peak isometric force (*P*_*0*_) was the mean force during the tetanus preceding ramp measurements. Force (*P*) was the force at the end of the ramp (**Figure 1**). Forces were converted to stresses by dividing by the CSA.

#### Power output

The power output of muscle was estimated by calculating the product of stress*strain from the force-velocity curves. To quantify the degree of curvature of force velocity curves, we calculated the power ratio as *Π*_*i*_ ÷ (*σ*_0_ × *V*_*max*_), where *Π*_*i*_ is the peak instantaneous power. Power ratios range from 0 to 1 with lower values indicating a higher degree of curvature.

#### Performance extrapolation to body temperature

Unfortunately, we were unable to effectively measure force-velocity at physiological body temperature of 39-41°C (Ellerby and Askew, 2007) due to the muscle shortening too rapidly for the high-speed length controller. Therefore, we instead measured at 20 and 30°C, and extrapolated to a physiological temperature of 40°C using *Q*_*10*_ values:

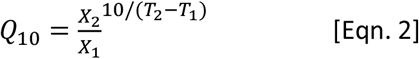

where *X*_*2*_ is velocity or other parameter at the higher temperature (*T*_*2*_) and *X*_*1*_ is the same parameter at the lower temperature (*T*_*1*_). These values were subsequently used to estimate the maximum shortening velocity at physiological temperature of 40°C.

#### Isometric contractile performance

To assess if previous measures of isometric contractile function are related to non-isometric properties we assessed four measures of isometric performance: the time to peak twitch force (*tP*_*tw*_), the time to peak tetanus force (*tP*_*0*_), the full-width of the twitch at half force (FWHF) and the twitch half relaxation time (*RT*_*50*_). All isometric contractile performance measures were assessed at *L*_*0*_ and 30°C.

### Statistical analysis

Statistical testing was conducted using IBM SPSS Statistics 28 (International Business Machines Corporation, Armonk, NY, USA). Data are presented as mean ± standard deviation (S.D.) throughout. All data were tested for normality (Shapiro-Wilk test) and homogeneity (Levene’s test) prior to statistical testing. Data which were non-normally distributed were transformed to meet these assumptions *via* log transformation. Data which could not be transformed, or where transformation was inappropriate, were tested for statistical differences using nonparametric tests. A critical p value of 0.05 was used to indicate significant differences. Male-female comparisons were made using independent samples t-tests or Mann-Whitney U tests. Comparisons of force-length methodologies were made using paired t-tests. Correlations between isometric contractile dynamics and shortening velocities were made using Pearson’s correlation coefficient.

## Results

### Force velocity relationship of syringeal muscle is not sex dependent

We successfully measured force-velocity curves in superfast zebra finch syringeal muscles using the isovelocity protocol. We used *Q*_*10*_ values to extrapolate DTB function at 40°C from measures taken at 20 and 30°C. In zebra finch syringeal muscle greater muscular force is associated with decreased velocities, typical for skeletal muscle (**Figures 1 & 2**). Previous work has found male-female differences in the twitch dynamics and muscle fibre types of zebra finches. Here we hypothesised male-female differences would also be present in the non-isometric properties of muscle function. We found male muscle *V*_*max*_ was 17% faster than female (**Table 1**), with power being on average 56% higher (**Table 1**), however these performance differences were not significant **(Table 1**; **Figure 3**). We also found the velocity at maximum power, relative stress at maximum power and power ratios were not different between male and female muscles. Taken together, these findings indicate force-velocity curves do not differ significantly between male or female muscle at physiological temperatures and we pooled the data. At 40°C, we estimated a peak shortening velocity of 42.87 ± 17.53 L_0_ s^-1^ (N = 14), with a power output of 36.51 ± 29.84 mW g^-1^. This power output was achieved at 36 ± 21% of *V*_*max*_ and 37 ± 28% of *σ*_*0*_. The power ratio of force-velocity curves at 40°C was 0.13 ± 0.11, indicating a low degree of curvature.

**Figure 2:**
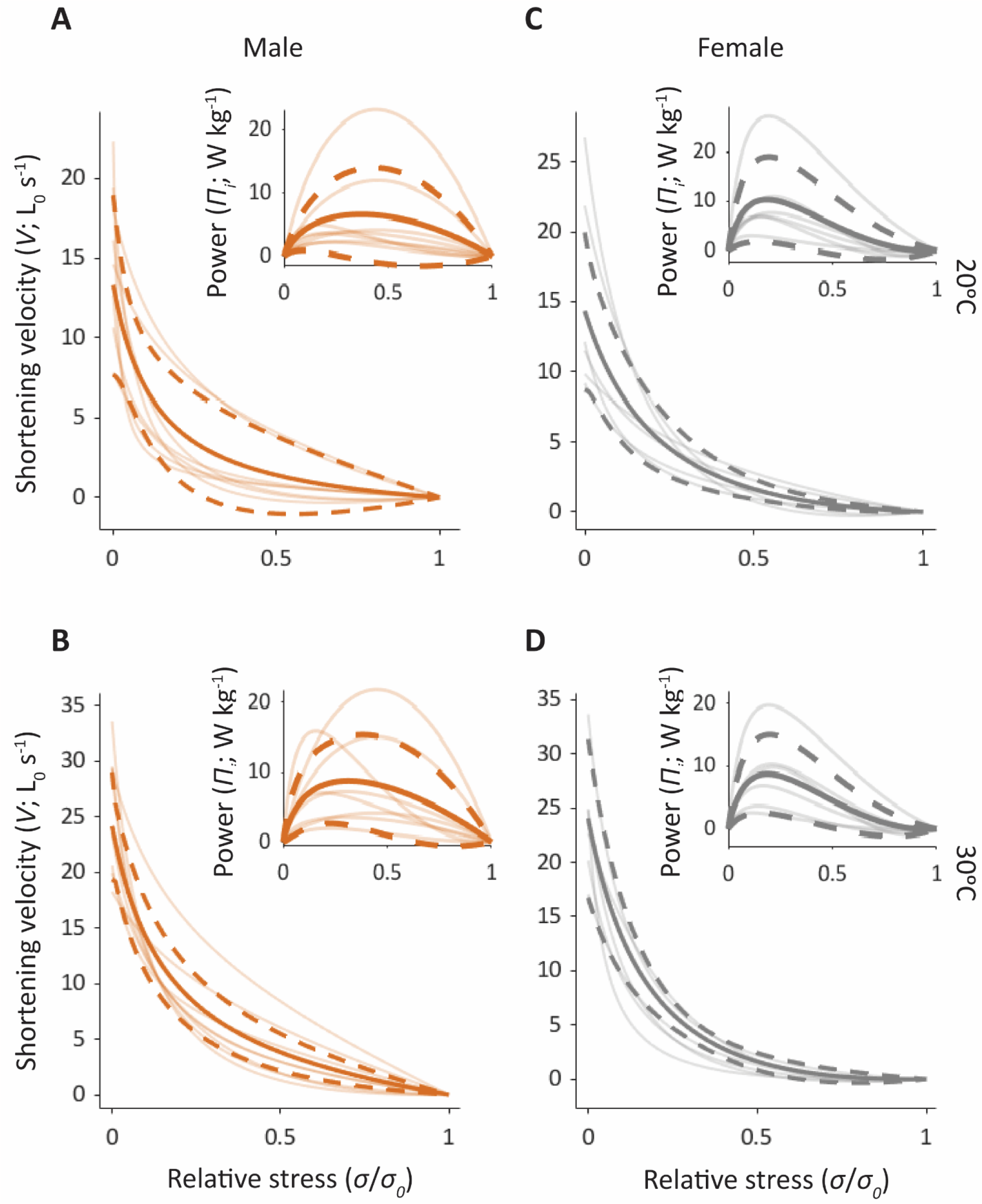
Force-velocity and force-power relationships in zebra finch syrinx muscle. A) Force-velocity curves of male (orange; N=8) DTB muscle at 20°C and (B) 30°C. (C) Force-velocity curves of female (grey; N=6) DTB at 20°C and (D) 30°C. (A-D) Insets show the resulting force-power relationships, calculated as stress*strain. The mean (solid) and standard deviation (dashed) are thick lines while individual curves are translucent.

**Figure 3:**
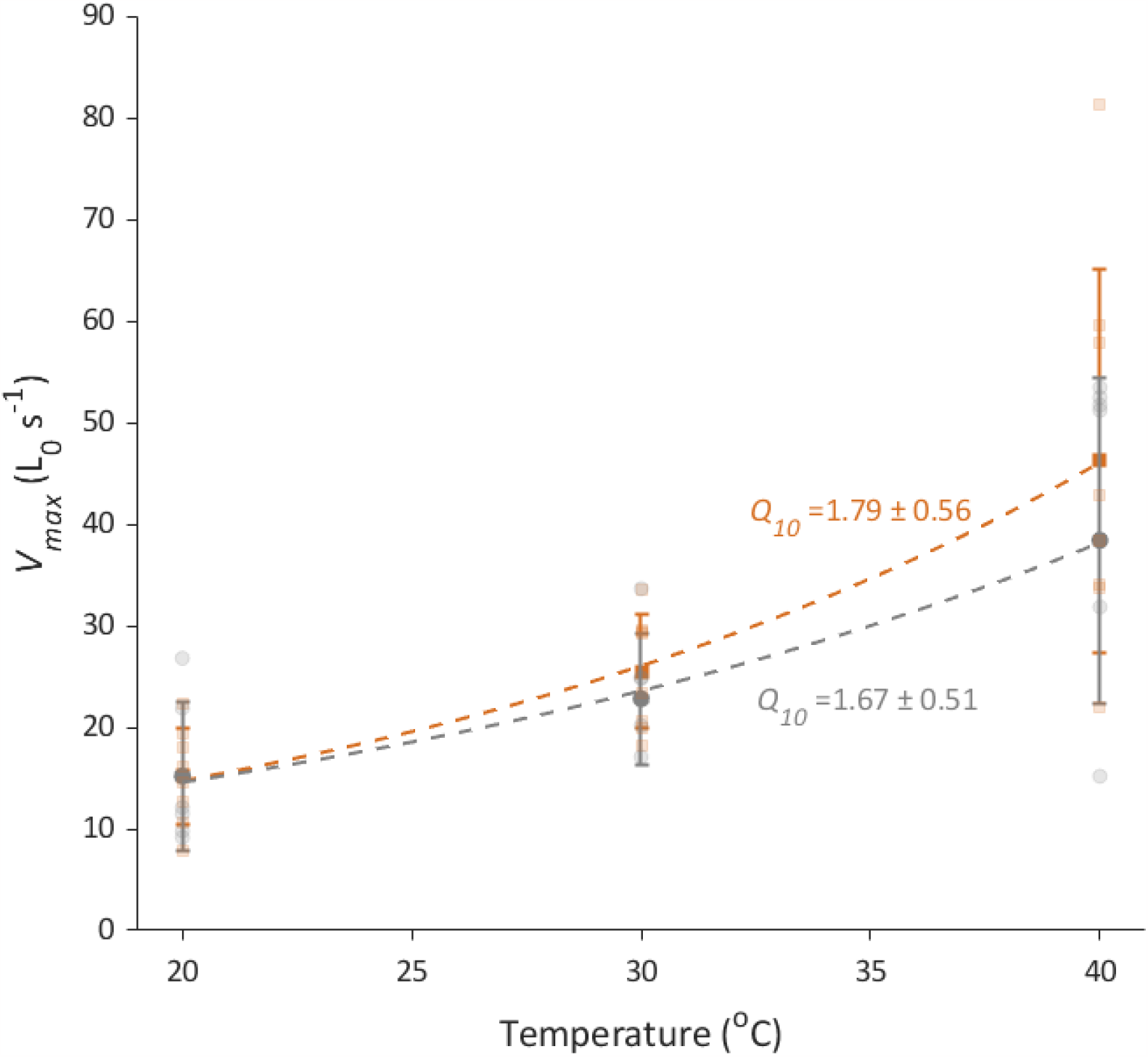
Maximum shortening velocity extrapolation of DTB muscle to body temperature. Measures were taken at 20 and 30°C in male (orange; N=8) and female (grey; N=6) DTB muscle and shortening velocity at 40°C estimated from these measures. Mean ± S.D. are shown as filled squares and circles; raw data are shown as translucent. The mean ± S.D. *Q*_*10*_ temperature coefficients are shown for each sex. With increased temperature, male and female muscle *V*_*max*_ starts to separate, variability is however high.

**Table 1:** Muscle properties of the DTB muscle of the zebra finch syrinx. Data are shown for the population mean (N=14), male (N=8) and female (N=6) animals at 20 and 30°C. 40°C measures are estimated using *Q*_*10*_ values. Statistical comparisons between male and female muscle are shown, with any significant differences (p *≤*0.05) highlighted in bold.

|  | Temperature (°C) | Population mean | Female | Male | Statistics |
| --- | --- | --- | --- | --- | --- |
| $\sigma_0$ (mN mm<sup>-2</sup>) | 20 | | **4.68 ± 1.82** | **2.47 ± 1.28** | **t = -2.97, df = 12, p = 0.012** |
|  | 30 |  | **7.05 ± 2.76** | **4.45 ± 1.81** | **t = -2.49, df = 12, p = 0.028** |
|  | Estimated 40 |  | **9.64 ± 5.03** | **6.78 ± 3.15** | **t = -2.42, df = 12, p = 0.032** |
| $\sigma$ at $\Pi_i$ ($\sigma/\sigma_0$) | 20 | 0.32 ± 0.08 | 0.35 ± 0.07 | 0.31 ± 0.09 | U = 31, n = 14, p = 0.41 |
|  | 30 | 0.31 ± 0.09 | 0.32 ± 0.09 | 0.31 ± 0.09 | U = 26, n = 14, p = 0.85 |
|  | Estimated 40 | 0.37 ± 0.28 | 0.35 ± 0.25 | 0.39 ± 0.32 | t = 0.07, df = 12, p = 0.95 |
| $V_{max}$ (L<sub>0</sub> s<sup>-1</sup>) | 20 | 15.20 ± 5.74 | 15.21 ± 7.32 | 15.20 ± 4.80 | t = 0.18, df = 12, p = 0.86 |
|  | 30 | 24.34 ± 5.91 | 22.81 ± 6.45 | 25.49 ± 5.63 | t = 0.91, df = 12, p = 0.38 |
|  | Estimated 40 | 42.87 ± 17.53 | 38.40 ± 15.99 | 46.22 ± 18.93 | t = 0.84, df = 12, p = 0.42 |
| $V$ at $\Pi_i$ ($V/V_{max}$) | 20 | 0.29 ± 0.09 | 0.31 ± 0.05 | 0.28 ± 0.11 | t = -0.61, df = 12, p = 0.55 |
|  | 30 | 0.31 ± 0.08 | 0.30 ± 0.06 | 0.31 ± 0.09 | t = 0.29, df = 12, p = 0.78 |
|  | Estimated 40 | 0.36 ± 0.21 | 0.30 ± 0.12 | 0.41 ± 0.26 | t = 0.41, df = 12, p = 0.69 |
| $\Pi_i$ (mW g<sup>-1</sup>) | 20 | | **15.50 ± 17.22** | **4.14 ± 4.84** | **t = -2.59, df = 12, p = 0.024** |
|  | 30 | 11.77 ± 6.69 | 12.87 ± 4.62 | 10.96 ± 8.13 | t = -1.06, df = 12, p = 0.31 |
|  | Estimated 40 | 36.51 ± 29.84 | 21.29 ± 7.65 | 47.93 ± 35.56 | t = 1.38, df = 12, p = 0.19 |
| Power ratio<br>($\Pi_i/\sigma_0 * V_{max}$) | 20 | 0.10 ± 0.04 | 0.11 ± 0.04 | 0.09 ± 0.04 | U = 32, n = 14, p = 0.35 |
|  | 30 | 0.10 ± 0.03 | 0.10 ± 0.04 | 0.09 ± 0.03 | U = 26, n = 14, p = 0.85 |
|  | Estimated 40 | 0.13 ± 0.11 | 0.12 ± 0.13 | 0.14 ± 0.10 | U = 20, n = 14, p = 0.66 |

Abbreviations: $\sigma_0$ peak isometric stress, $\sigma$ stress, $V_{max}$ maximum velocity of muscle shortening at zero force, $V$ muscle shortening velocity, $\Pi_i$ instantaneous power.

The influence of temperature on DTB muscle performance was greater in male than female muscle. At 20°C, male muscle had significantly lower power (*t* = -2.59, df = 12, p = 0.024; **Table 1; Figure 2**). These differences in thermal dependence are reflected in *Q*_*10*_ values (Table 2), where muscle power output has a *Q*_*10*_ value of 4.37 ± 3.85 in males vs 1.65 ± 1.49 in females.

**Table 2:** Temperature coefficients of measured parameters. Data are shown for the population mean (N=14), male (N=8) and female (N=6) muscles. Temperature coefficients are calculated from measures at 20 and 30°C.

|  | Population mean | Female | Male |
| --- | --- | --- | --- |
| $\sigma_0$ | 1.48 ± 0.37 | 1.33 ± 0.16 | 1.58 ± 0.17 |
| $\sigma$ at $\Pi_i$ | 1.07 ± 0.54 | 1.00 ± 0.48 | 1.13 ± 0.61 |
| $V_{max}$ | 1.74 ± 0.52 | 1.67 ± 0.51 | 1.79 ± 0.56 |
| $V$ at $\Pi_i$ | 1.10 ± 0.43 | 0.99 ± 0.21 | 1.19 ± 0.54 |
| $\Pi_i$ | 3.21 ± 3.29 | 1.65 ± 1.49 | 4.37 ± 3.85 |
| Power ratio<br>($\Pi_i/\sigma_0 * V_{max}$) | 1.23 ± 0.78 | 1.04 ± 0.73 | 1.37 ± 0.83 |

Abbreviations: $\sigma_0$ peak isometric stress, $\sigma$ stress, $V_{max}$ maximum velocity of muscle shortening at zero force, $V$ muscle shortening velocity, $\Pi_i$ instantaneous power.

### Relationship between isometric contractile dynamics and shortening velocity

The twitches of male muscles were significantly faster than female (**Figure 4**), with full width at half maximal force (FWHF) times of 9.79 ± 5.15 ms in males and 14.48 ± 4.10 ms in females at 30 °C (*t =* -0.21, df = 12, p = 0.031). Other aspects of twitch performance, the time to peak twitch (*tP*_*tw*_) and twitch half relaxation time (*RT*_*50*_), were also significantly faster in male than female muscle, where *tP*_*tw*_ was 5.69 ± 1.27 ms in males and 9.14 ± 2.17 ms in females (*t =* -3.62, df = 12, p = 0.004). *RT*_*50*_ was 5.50 ± 3.13 ms in males and 8.55 ± 2.59 ms in females (*t =* -2.25, df = 12, p = 0.044). However, the time to peak tetanus (*tP*_*0*_) was not significantly different between sexes (*t =* -1.98, df = 12, p = 0.072; male DTB: 38.90 ± 16.00 ms, female DTB: 55.63 ± 16.74 ms) and was 46.07 ± 17.87 (N=14).

**Figure 4:**
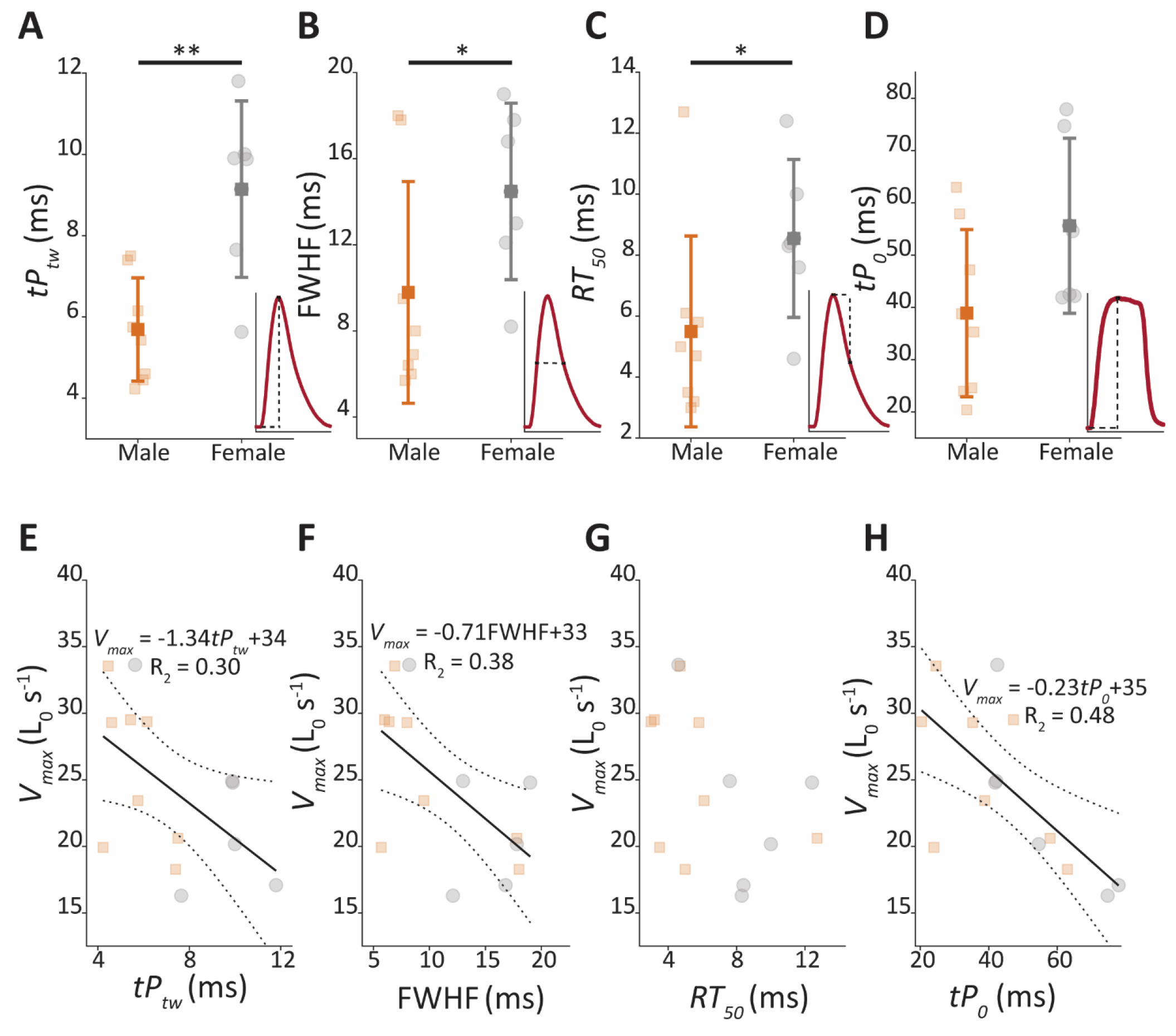
Isometric properties that predict muscle shortening velocity. Isometric contractile features of DTB muscle at 30°C: (A) The time to peak twitch (*tPtw), (*B) full width at half maximal force time (FWHF), (C) twitch half relaxation time (*RT*_*50*_) and (D) time to peak tetanus (*tP*_*0*_). Superimposed panels in (A) – (D) give examples of how these parameters were measured. (E) The relationship between *V*_*max*,_ and *tP*_*tw*_, (F) FWHF, (G) *RT*_*50*,_ and (H) *tP*_*0*_. Correlation equations shown in (E), (F) & (H) were all significant. Data from males (N=8) is shown in orange, and females (N=6) in grey, translucent colours show individual datapoints, solid colours show the mean ± S.D.

To assess if commonly used isometric contractile parameters can predict *V*_*max*_, we assessed if any of the measured properties correlated. Of the measured twitch parameters, we found both *tP*_*tw*_ and FWHF negatively correlated with *V*_*max*_ (*tP*_*tw &*_ *V*_*max*_ r= -0.55, N = 14, p = 0.044; FWHF & *V*_*max*_ r = -0.68, N = 14, p = 0.018), but *RT*_*50*_ did not correlate significantly with *V*_*max*_ (r = -0.47, N = 14, p = 0.093). Of the measured tetanic parameters, the time to peak tetanus, *tP*_*0*_ negatively correlated with *V*_*max*_ (r = -0.69, N = 14, p = 0.006). Faster twitches and tetani are thus associated with increased shortening speed (**Figure 4**).

### Comparison of force-length methodologies

Previously, laryngeal muscle force-length properties have been determined through a cyclic methodology (Alipour-Haghighi et al., 1991; Alipour and Titze, 1999). Lastly, we tested if this methodology provides similar force-length profiles as a stepwise methodology for syringeal muscles (**Figure 5**). Stepwise methodologies gave mean *L*_*0*_ values of 5.16 ± 0.23 mm and 4.98 ± 0.23 mm in males and females respectively. Cyclic methodologies gave mean *L*_*0*_ values of 5.07 ± 0.23 mm and 5.06 ± 0.24 mm. Peak stresses at *L*_*0*_ were 4.45 ± 0.64 mN mm^-2^ in males and 7.05 ± 1.13 mN mm^-2^ in females using stepwise methodologies, while cyclic methodologies returned peak stresses of 3.60 ± 0.72 mN mm^-2^ in males and 5.42 ± 0.64 mN mm^-2^ in females (**Figure 5F**). Pairwise comparisons revealed methodologies did not significantly differ in *L*_*0*_ (*t* = 0.40, df = 13, p = 0.70) or *σ*_*0*_ (*t* = 0.25, df = 13, p = 0.81). Thus, both methodologies result in the same values for *L*_*0*_ in both male and female preparations (**Figure 5E**).

**Figure 5:**
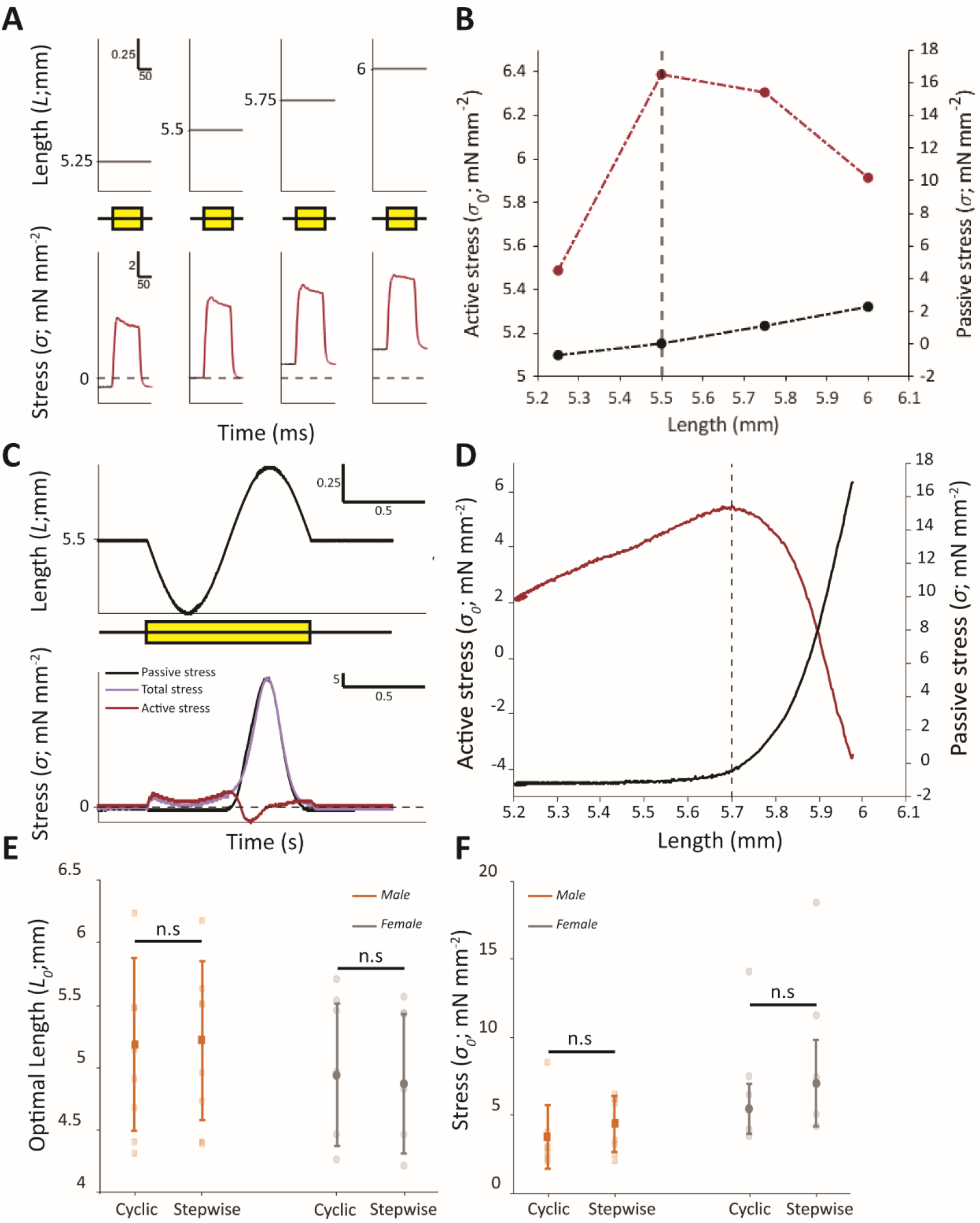
Cyclic and stepwise protocols provide the same optimal length and stress in vocal muscles. (A) stepwise approach muscle length is increased in steps and tetanised at each length. (B) Resulting profile from this approach with active force shown in red, and passive in black, vertical dashed line shows the *L*_*0*_ length. (C) Cyclic approach, muscle is stimulated throughout a 1 Hz length change (± 0.5 mm from starting length), top panel shows length change, and bottom shows resulting stress outputs. Black line shows the passive stress, purple total stress and red active stress (calculated as total stress minus passive stress). (D) Resulting profile from approach (C) active stress shown in red and passive in black, the vertical dashed line indicates *L*_*0*._ Yellow boxes in (A & C) are used to show the stimulation period. (E) The optimal length (*L*_*0*_) from the stepwise and cyclic methodologies. Solid filled orange squares (male; N=8) and grey circles (female; N=6) show the mean ± S.D., translucent squares and circles show raw data. (F) The stress (*σ*_*0*_) at the optimal length during stepwise and cyclic methodologies. Solid filled orange squares (male) and grey circles (female) show the mean ± S.D., translucent squares and circles show raw data.

Finally, to test if the prolonged stimulation (1 minute) during cyclic approaches impacted muscle function, we tested if tetani 5 minutes after this, differed from those before. We found no significant differences in male (before: 4.11 ± 2.13 mN mm^-2^, after: 3.44 ± 0.92 mN mm^-2^; *t =* 0.54, df = 7, p = 0.61) or female (before: 5.53 ± 2.57 mN mm^-2^, after: 6.38 ± 3.15 mN mm^-2^; *t* = -1.06, df = 5, p = 0.34) DTB muscle (**Figure 6**).

**Figure 6:**
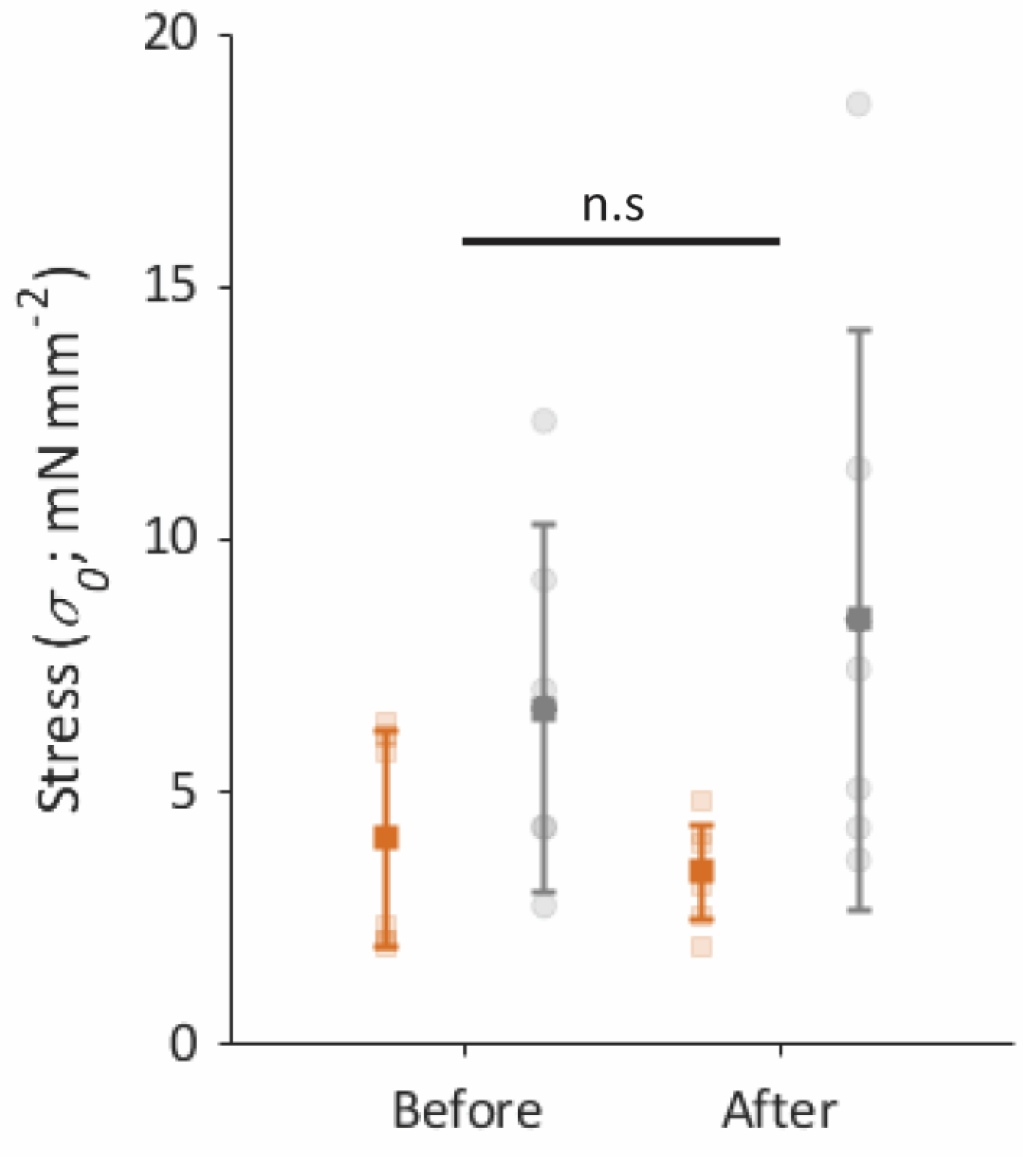
Prolonged stimulation does not impair muscle function. Isometric stress before and after being stimulated for 1 minute. Stress 5 minutes after prolonged stimulation did not significantly differ from previous tetani. Male data (N=8) are shown in orange, and female (N=6) in grey, solid colours show the mean ± S.D., translucent show individual data points.

## Discussion

With a maximum shortening velocity of 46 L_0_ s^-1^, and one individual up to 81 L_0_ s^-1^, at body temperature, zebra finch superfast syringeal muscle has the highest measured shortening velocity to date. It far exceeds the shortening velocity seen in other avian fast-twitch muscles, such as cockatiel (*Nymphicus hollandicus*), zebra finch and blue-breasted quail (*Coturnix chinensis*) pectoralis muscle, with respective maximum velocities of 15, 21 and 32 L_0_ s^-1^ (Askew and Marsh, 2001; Ellerby and Askew, 2007; Morris and Askew, 2010). Compared to other established superfast muscles at physiologically relevant temperatures, syringeal muscle has substantially faster shortening velocity. Rattlesnake (*Crotalus atrox*) shaker muscle reaches only 18 L_0_ s^-1^ at 35°C (Rome and Lindstedt, 1998; Rome et al., 1996). We currently lack force-velocity data on toadfish swimbladder and bat laryngeal at physiological temperatures (Elemans et al., 2011; Rome, 2006). Rabbit extraocular muscle, a potential superfast muscle, shortens at 23 L_0_ s^-1^ at 35°C (Briggs and Schachat, 2000). Comparisons with other vocal muscles, reveal considerably faster shortening in syringeal muscle. The thyroarytenoid (TA), a laryngeal muscle, of humans (*Homo sapiens*) has a *V*_*max*_ between 2.55 (D’Antona et al., 2002) and 2.9 L_0_ s^-1^ (Sciote et al., 2002) at 12 and 15°C. In baboons, TA shortens up to 6.7 L_0_ s^-1^ at body temperatures (37°C) (Mardini et al., 1987). TA muscle achieved shortening velocities of up to 13 L_0_ s^-1^ at 25°C in rat, and < 2.68 L_0_ s^-1^ at 12°C (Toniolo et al., 2007), and 6.7 L_0_ s^-1^ at a body temperature of 37°C in dogs (Alipour and Titze, 1999). The high shortening velocities reported here are, to our best knowledge, only matched by values estimated for myotomal muscles in larval zebrafish (up to 45 L_0_ s^-1^ at 28 °C) (Mead et al., 2020) that are used for fast C-starts in escape responses. The differences in maximal shortening velocity between muscles reflects their optimisation for different features and different functions. The extremely high shortening velocities in vocal muscles again highlights they are tuned for speed.

Our data show that fast isometric properties correlate positively with maximal shortening velocities, corroborating previous work (Close, 1965; Lännergren, 1978). Despite these correlations, and even though our isometric measures confirm male twitches are nearly twice as fast as female twitches (Adam and Elemans, 2020), superfast shortening was not significantly different between sexes. Closer examination of the data presented here reveals that the *V*_*max*_ distributions of male and female muscle is starting to separate at 40°C (**Figure 3**). Therefore, we suspect that with additional birds, data distribution would separate more strongly. Additionally, disparities between twitch speed and shortening velocities have been shown in other muscles (Gladman and Askew, 2022; Marsh, 1990). For example, in cuttlefish (*Sepia officinalis*) mantle muscle, juveniles have significantly faster twitches than adults, but shortening velocities do not differ between the two age groups (Gladman and Askew, 2022). These disparities were suggested to result from twitch dynamics being more closely aligned with cyclic muscle performance than with shortening velocity (Gladman and Askew, 2022; Marsh, 1990). Twitch activation and relaxation times are determined by the deactivation of crossbridge cycling. The high energetic costs associated with rapid deactivation mean that isometric contractile properties are usually tightly coupled with the operating frequency of muscle seen during movement and, in this case, song. This is achieved by maintaining a specific ratio between contraction kinetics and cycle duration, regardless if the muscle is activated through a single stimulus or through a burst of stimuli (Askew and Marsh, 2001; Gladman and Askew, 2022; Marsh, 1990). Recent work may confirm this hypothesis in syringeal tissues, where disuse of syringeal muscles leads to muscles becoming significantly slower, and fibre types shifting from superfast to fast (Adam et al., 2023). These changes are coupled with a reduction in parvalbumins, calcium handling systems associated with the deactivation of crossbridge cycling. The reduced deactivation capacity and slowing of twitch dynamics, suggests high deactivation rates are required by syringeal muscle to achieve cycle frequencies ≥200 Hz. If twitch dynamics are strongly coupled with cyclic performance in zebra finch syringeal muscle, we expect female muscles will operate at lower cycle frequencies, but we currently lack data to empirically test this hypothesis.

Our data show that conventional stepwise approaches and continual stimulation during cyclic motion previously employed in laryngeal tissues (Alipour-Haghighi *et al*. 1991) provides the same *L*_*0*_ and stress. During cyclic continual stimulation passive stress is higher beyond *L*_*0*_, likely because the muscle is not reaching a steady state through the experimental procedure. Surprisingly, however, muscles subject to continual stimulation did not show significant changes in muscle force output, suggesting no irreversible damage to the tissue occurred. Stepwise approaches take substantially more time to build force-length profiles, cyclic approaches can find *L*_*0*_ in under two seconds, while stepwise approaches typically require 15 minutes, over which time muscle may degrade. Therefore, the cyclic approach could act as a means to get the most from the tissue before it degrades further. Taken together, we suggest either method is appropriate for finding force-length properties of vocal muscles, but we recommend to use the stepwise method where measurements of passive stress better reflect the steady-state.

In tuning syringeal muscle to operate at high frequencies, we show that next to force, instantaneous power has also been traded for speed. In zebra finches the force output of syringeal muscle is 95-96% lower than that of the fast-twitch pectoralis (Ellerby and Askew, 2007). Reduced force generating capacity results from decreased myofibrillar area (Mead et al., 2017), and crossbridge duty cycles (Askew, 2023), which together with high detachment rates enable rapid shortening and cycling rates (Rome et al., 1999; Tikunov and Rome, 2009). The reduction in force-generating capacity is not adequately compensated for by increased speed, resulting in instantaneous power of syringeal muscle being substantially lower than pectoralis muscle (Ellerby and Askew, 2007). Instantaneous power is much greater than cyclic power. Elemans *et al*. (2008) reported cyclic power reaches ∼6 W kg^-1^ at frequencies <100 Hz. At higher frequencies (200 Hz) this declines to ∼2 W kg^-1^. Disparities between cyclic and instantaneous power is a common feature, where cyclic power output is between 15-33% of instantaneous power in locomotory muscles (Ellerby and Askew, 2007; Gladman and Askew, 2022; James et al., 1996). We estimate the maximum cyclic power output is 13-29% of the instantaneous power output, which agrees with in locomotory muscles, and suggests the underlying mechanical processes are similar between muscle types.

Superfast syringeal muscles performed too fast to measure their FV directly at body temperature using the isovelocity technique. Therefore, we estimated force-velocity performance at body temperate by extrapolation using *Q*_*10*_ thermal dependence, these values were similar to reported values. *V*_*max*_ *Q*_*10*_ values of rabbit extraocular muscle and zebrafish myotomal muscle are 1.7 and 2.2 (Asmussen et al., 1994; Mead et al., 2020), and the DTB muscle falls within this range (*V*_*max*_ *=* 1.74). Wider comparisons reveal thermal-dependence of rate-based processes is a common feature of muscle (see Bennett, 1984 & 1985). Shortening velocity of locomotory muscle has a typical *Q*_*10*_ value of between 1.4 and 2.2. Thus, the thermal dependence *Q*_*10*_ and therefore reported *V*_*max*_ values fall within the range of previously reported values and seem realistic.

The high thermal dependence of superfast muscles we demonstrated suggests that muscle function is compromised at non-physiological temperatures. This is also shown e.g in hummingbird pectoralis muscle., where drastic temperature reductions of *≤*20°C produce substantially reduced force (Reiser et al., 2013). Even in endotherms, such significant reduction of body temperature are commonly seen and behaviourally relevant: Small passerines reduce body temperature 5-10°C during the winter (Brodin et al., 2017; García-Díaz et al., 2023; Ruf and Geiser, 2015) and hummingbirds and bats reduce body temperature *≤*20°C during daily torpor to reduce energetic costs (Luo et al., 2021; Shankar et al., 2022). The underlying muscle mechanics may now affect behavioural performance. Indeed Wu *et al*. (2021) found the echolocation call frequency of leaf-nosed bats (*Hipposideros armiger*) decreased with decreased body temperature, and great tits (*Parus major*) switch from defending territories through elaborate song to using short, less demanding, alarm calls during cold days (Strauss et al., 2020). Thus, the thermal dependence in superfast vocal muscle may impact vocal behaviour.

## Acknowledgements

We thank Timothy G. West for supplying the T-clips, and Maria Anthonsen for technical assistance. We also thank Sonja Jacobsen, Emilie Radoor and Emilie Jensen for animal care and maintenance.

## Data availability

Data are available from figshare: [these will be upload and a DOI provided prior to publication]

---

**Supplementary Table 1:**
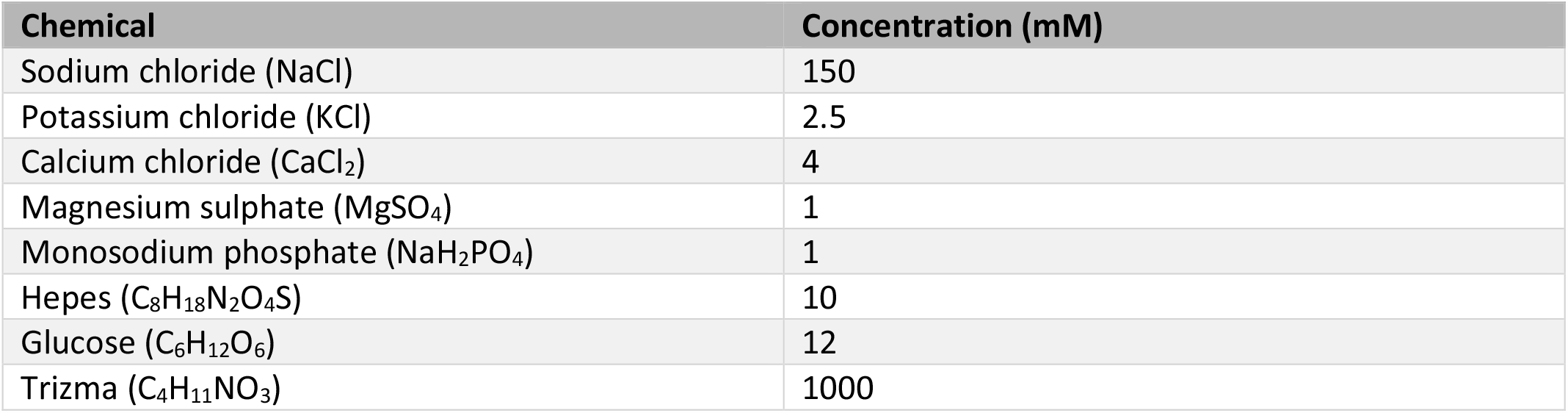
Chemical composition of avian ringers solution. Solution was adjusted to pH 7.4 using Trizma.

